# Unraveling nitrogen, sulfur and carbon metabolic pathways and microbial community transcriptional responses to substrate deprivation and toxicity stresses in a bioreactor mimicking anoxic brackish coastal sediment conditions

**DOI:** 10.1101/2021.08.31.458400

**Authors:** Paula Dalcin Martins, Maider J. Echeveste, Arslan Arshad, Julia Kurth, Heleen Ouboter, Mike S. M. Jetten, Cornelia U. Welte

## Abstract

Microbial communities are key drivers of carbon, sulfur and nitrogen cycling in coastal ecosystems, where they are subjected to dynamic shifts in substrate availability and exposure to toxic compounds. However, how these shifts affect microbial interactions and function is poorly understood. Unraveling such microbial community responses is key to understand their environmental distribution and resilience under current and future disturbances. Here, we used metagenomics and metatranscriptomics to investigate microbial community structure and transcriptional responses to prolonged ammonium deprivation and sulfide and nitric oxide toxicity stresses in a controlled bioreactor system mimicking coastal sediment conditions. *Candidatus* Nitrobium versatile, identified in this study as a sulfide-oxidizing denitrifier, became a rare community member upon ammonium removal. The methanotroph *Ca.* Methanoperedens nitroreducens showed remarkable resilience to both experimental conditions, dominating transcriptional activity of dissimilatory nitrate reduction to ammonium (DNRA). After the ammonium removal experiment, a novel methanotroph species that we have named *Ca.* Methylomirabilis tolerans outcompeted *Ca.* Methylomirabilis lanthanidiphila and the anaerobic ammonium oxidizer (anammox) *Ca.* Kuenenia stuttgartiensis outcompeted *Ca.* Scalindua rubra. At the end of the sulfide and nitric oxide experiment, a gammaproteobacterium affiliated to the family *Thiohalobacteraceae* was enriched and dominated transcriptional activity of sulfide:quinone oxidoreductase. Our results indicate that some community members could be more resilient to stresses than others in coastal ecosystems, leading to dynamic microbial community shifts and novel functional states. Methane and sulfide oxidation could be ecosystem functions preserved across the investigated disturbances, while differing nitrogen cycling pathways might be favored in response to stresses.

**Importance:** Coastal ecosystems are primary zones of biogeochemical cycling, processing inputs of nutrients both generated *in situ* and derived from land runoff. Microbial communities that inhabit costal sediments perform these biogeochemical reactions, but microbial responses to dynamic, periodic substrate deprivation and exposure to toxic compounds remain elusive. In this study, we sought to address this knowledge gap in a controlled bioreactor system, unraveling microbial metabolic pathways and monitoring microbial responses to stresses that might occur in costal sediments. We identified key microbial players and shifts in their abundance and transcriptional activity. Our results indicated that methanotrophs were particularly resilient to stresses, sulfide oxidizers differed in resiliency but the community maintained sulfide oxidation function across stresses, and that anaerobic ammonium oxidizing (anammox) bacteria were sensitive to substrate deprivation but could restore activity once favorable conditions were reestablished. These insights will help to understand and predict coastal ecosystem responses to future disturbances.

## Introduction

Microorganisms drive and link the biogeochemical cycles of carbon, nitrogen and sulfur by a variety of redox reactions (1). Anthropogenic nutrient inputs from land into the ocean constitute a major impact on marine ecosystems, altering seawater and sediment biogeochemistry and leading to increased primary production that can result in toxic algal blooms and oxygen depletion (2). Such impacts, combined with ocean warming and consequent seawater stratification and deoxygenation, can further stimulate the production of methane and nitrous oxide, potent greenhouse gases, as well as sulfide and nitric oxide, toxic products of sulfate reduction and denitrification (2–5). In coastal sediments, ammonium and nitrate can be introduced via agricultural runoff, while sulfide, nitrogen oxides, methane, and ammonium are generated *in situ* via sulfate reduction, partial denitrification, methanogenesis, and organic matter decomposition, respectively (6). Therefore, characterizing microbial communities, interactions and reactions performed by microorganisms that couple methane, nitrogen and sulfur cycling is fundamental for understanding biogeochemical cycling and linked greenhouse gas emissions in dynamic coastal ecosystems impacted by anthropogenic activity.

Modeling efforts suggest links between future environmental changes, biogeochemical cycles and ecosystem functions (7, 8). However, given that most microorganisms are widespread and functionally redundant, they are frequently treated as a “black box” in models - preventing the effective modeling of their reactions and responses. Additionally, recent efforts with laboratory cultures and engineered systems have obtained insights about impacts of substrate availability changes on environmentally and economically relevant microbial communities (9–14). Yet, few studies have examined microbial community responses to prolonged periods of substrate scarcity or environmental stresses in controlled systems (15, 16). However, these are highly relevant disturbances in coastal ecosystems, where, for instance, nitrogen limitation is a major control on eutrophication (17) and sulfide toxicity can lead to mortality of marine life (18). Such studies are needed to unravel key microbial interdependencies, competitive interactions and functional shifts, as well as to comprehend their environmental distribution and resilience under current and future disturbances.

Carbon-, nitrogen- and sulfur-cycling microbial communities harbor potential for biotechnological applications such as the improvement of wastewater treatment systems. For instance, Deng *et al.* proposed that sulfide-driven partial denitrification could be coupled to anaerobic ammonium oxidation (anammox) in future applications, given that rapid oxidation of sulfide to elemental sulfur can prevent toxicity and inhibition of anammox activity (13). On the other hand, sulfide addition in a controlled aerated bioreactor setting promoted undesirable production of nitrous oxide and nitrite via partial denitrification and dissimilatory nitrite reduction to ammonium (DNRA), respectively (14). These observations indicate that further studies are necessary to understand complex microbial community interactions and to evaluate how they can be best employed in biotechnological applications.

In this research project, we investigated transcriptional stress responses of a complex microbial community enriched in an anoxic laboratory-scale bioreactor mimicking dynamic, brackish sediment conditions, where periodic ammonium deprivation, and sulfide and nitric oxide (NO) toxicity stresses, the chosen stressors in this study, might occur. The culture performed sulfide, ammonium and methane oxidation at the expense of nitrate via sulfide-oxidizing denitrifiers, anammox bacteria, and nitrite/nitrate-dependent anaerobic methane oxidizers (19). This study’s aims were (1) to understand the effect of periodic ammonium removal on dissimilatory nitrite reduction to ammonium (DNRA) as a source of ammonium for anammox activity, as well as on general microbial community structure and transcriptional activity, (2) to unravel potential metabolic reactions utilized by key community members, and (3) to characterize microbial community structure and transcriptional responses to prolonged sulfide and NO toxicity stresses while attempting to enrich sulfide-oxidizing denitrifiers.

## Materials and methods

### Bioreactor operation under regular and experimental conditions

A previously described complex co-culture of sulfide- and methane-oxidizing denitrifiers and anammox bacteria (19) was subjected to a 10-week substrate starvation (ammonium removal) experiment, and later, after regular operation conditions were re-established, to a 7-week sulfide and nitric oxide stress experiment (Figure 1). Briefly, under regular operation, the reactor culture was kept under anoxic conditions and fed daily with 0.67 mmol of sulfide, 1.4 mmol of ammonium, 3 mmol of nitrate, and 488 mmol of methane. The medium contained a solution of salts, trace elements, minerals and vitamins as previously described (19), and the pH was kept at 7.1. Under experimental operation, the following conditions were modified. During the ammonium removal experiment, the ammonium concentration in the medium was gradually decreased from 7 to 0 mM over one month. Two more weeks passed without ammonium addition to the reactor in order to ensure that no residual ammonium was present, and the first biomass sample for metatranscriptomic sequencing (T1) was collected and stored at −80°C. After 10 weeks more without ammonium, another biomass sample for metatranscriptomic sequencing was collected (T2) and ammonium was re-introduced into the medium.

**Figure 1.**
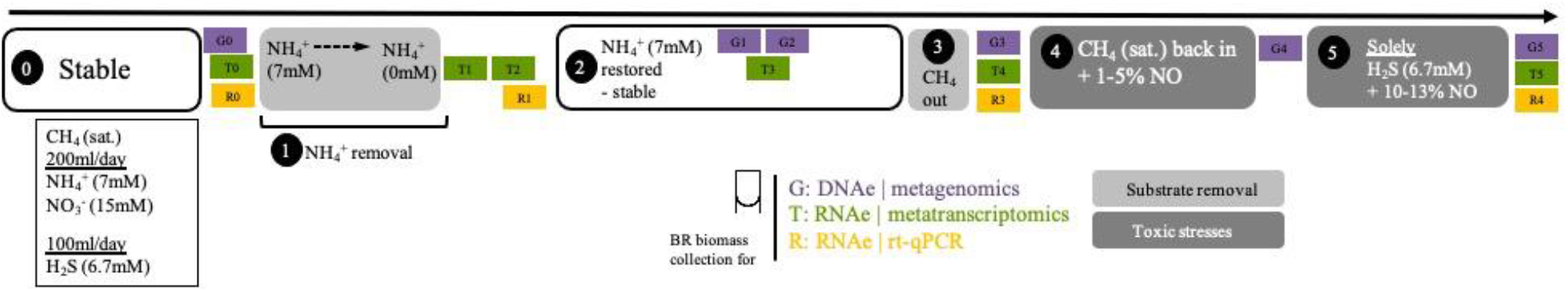
Timeline of experiments. Zero indicates when the bioreactor was stably maintained; 1, when the ammonium removal experiment happened; 2, after the ammonium removal experiment and re-stabilization period; 3; reactor placement under a fume hood in preparation for the next experiment; 4, preparatory period for the nitric oxide and sulfide toxicity stresses experiment; 5, nitric oxide and sulfide toxicity stresses experiment. Samples for metagenomics (G0-G5), metatranscriptomics (T0-T5) and RT-qPCR (R0-R1 and R3-R4) were collected as indicated across experimental time points and in between them. The dashed line in the ammonium removal experiment indicates that removal was conducted stepwise (in mM: 7 → 4.5 → 3.5 → 2.5 → 0). Exact dates and experiment durations are provided in Supplemental Table 1.

In preparation for the next experiment, the reactor was moved into a laminar flow hood. Methane flow was stopped at the time of reactor moving and reintroduced after 15 days. Once regular operation conditions and substrate consumption were reestablished, the sulfide and NO toxicity stress experiment began. Initially, the NO concentration in the reactor headspace was gradually increased to 1-5% for 10 weeks so that the community could adapt. Then, NO concentrations were increased to ~10-13% and all substrates except sulfide were removed from the reactor, resulting that only sulfide, NO and medium were provided to the reactor for the following 7 weeks. A biomass sample for metatranscriptomic sequencing was collected before any NO was added to the reactor (T4) and after the 17 weeks (T5). Originally, the experiment was planned to be conducted in 20 weeks, being that during the 10 last weeks the reactor would be maintained under only sulfide and NO. However, covid-19 lockdown in the Netherlands resulted in restricted access to the laboratory and the timing had to be changed. Additional metatranscriptomics samples in between experiments for were included in this study: T0, approximately 2 months before the ammonium removal experiment, and T3, 6 months after the ammonium removal experiment and 11 months before the sulfide and NO stress experiment (Figure 1).

During regular and experimental conditions, the reactor was checked daily for general parameters including pH with an Applisense electrode (Applikon, Delft, The Netherlands) and nitrate and nitrite concentrations with MQuantTM colorimetric test strips (Merck, Darmstadt, Germany). During experimental conditions, nitrate and nitrite were also measured with the Griess assay (Griess, 1879) as previously described (19), and ammonium was measured fluorometrically after reaction with 10% ortho-phthaldialdehyde as previously described (20). NO in headspace samples was measured with a Sievers Nitric Oxide analyzer (NOA280i; GE Power Water & Process Technologies, Boulder, CO), and sulfide in liquid samples was measured with the methylene blue assay (21) (HACH, Loveland, CO, USA). Sulfate was determined via the barium precipitation method using the Sulfate Assay Kit following manufacturer’s instructions (Sigma Aldrich, Saint Louis, MI, USA). Sample measurements were carried out in technical triplicates.

### Nucleic acid extractions and sequencing

Biomass samples for metagenomic sequencing were collected during regular and experimental periods (Figure 1) and stored at −20°C. All DNA extractions were performed using the DNAeasy Power Soil Kit (Qiagen, Hilden, Germany) following manufacturer’s instructions with two modifications: bead beating was performed in a TissueLyzer (Qiagen, Hilden, Germany) for 10 minutes and autoclaved MilliQ water was used instead of the kit’s buffer in the last elution step. RNA extractions were initially performed (for T1 and T2) using the RiboPure Bacteria Kit (Thermo Fisher Scientific, Waltham, MA, USA) following manufacturer’s instructions. During the stress experiment (T4 and T5), this method no longer resulted in sufficient extracted RNA for sequencing, potentially due to extensive extracellular polymeric substance production, and thus the method was changed to RNAeasy Power Soil Kit (Qiagen, Hilden, Germany), which was also used for two additional samples (T0 and T3). All RNA samples were treated with DNAase I at 37°C for 30 min from the RiboPure Bacteria Kit (Thermo Fisher Scientific, Waltham, MA, USA). DNA and RNA concentrations were measured with a Qubit 2.0 fluorometer using the dsDNA and RNA HS Kits (Thermo Fisher Scientific, Waltham, MA, USA). DNA and RNA quality were determined using a NanoDrop Spectrophotometer ND-1000 (Isogen Life Science, Utrecht, The Netherlands) and a Bioanalyzer 2100 (Agilent, Santa Clara, CA, USA), respectively.

Metagenomic libraries were prepared using the Nextera XT Kit (Illumina, San Diego, California, USA) following manufacturer’s instructions. Enzymatic tagmentation was performed with 1 ng DNA, followed by the incorporation of indexed adapters and amplification of the library. Amplified DNA libraries were then purified, and their quality and concentration were determined as aforementioned. Libraries were sequenced on an Illumina MiSeq platform (San Diego, California, USA) using the MiSeq Reagent Kit v3 (San Diego, California, USA), generating 300 bp paired-end reads. RNA samples of T1 and T2 were rRNA-depleted with MICROBexpress Kit (Thermofisher, Waltham, USA) and MEGAclear kit (Ambion, Life Technologie, Carlsbad, CA USA). Subsequently, 0.1 - 4 μg of RNA from T1-T3 were used to construct strand specific RNA-Seq libraries. Non-rRNA in RNA-Seq libraries were enriched by selective priming during the first strand cDNA synthesis reaction, as well as in the final library construction steps using TruSeq Stranded mRNA sample preparation guide (Illumina proprietary catalog RS-122-9004DOC). RNA from these samples was sequenced with an Illumina MiSeq platform (Illumina, CA, USA), generating 151 bp single-end reads. Metatranscriptomic libraries of T0, T4 and T5 were constructed using TruSeq stranded mRNA library Kit (Illumina, San Diego, California, USA). These RNA-Seq libraries were sequenced with an Illumina NovaSeq 6000 platform (San Diego, California, USA), generating 151 bp paired-end reads (San Diego, California, USA). All metatranscriptomes were triplicate RNA extractions and sequencing.

### Metagenomic and metatranscriptomic analyses

Metagenomic data were analyzed as it follows. Read quality was accessed with FASTQC v0.11.8 before and after quality-trimming, adapter removal and contaminant filtering, performed with BBDuk (BBTools v38.75). Trimmed reads were co-assembled *de novo* using MEGAHIT v1.2.911 (22) and mapped to assembled contigs using BBMap (BBTools v38.75) (23). Sequence mapping files were handled and converted using SAMtools v1.10. Contigs at least 1000 bp-long were used for binning with CONCOCT v1.1.0 (24), MaxBin2 v2.2.7 (25), and MetaBAT2 v2.1512 (26). Resulting metagenome-assembled genomes (MAGs) were dereplicated with DAS Tool v1.1.213 (27) and taxonomically classified with GTDB-Tk v1.3.0 (28) release 9514. MAG completeness and contamination was estimated with CheckM v1.1.2 (29).

MAGs were annotated with DRAM v1.0 (30)with default options, except -min_contig_size 1000 bp, and genes of interest were searched in annotation files as well as via BLASTP and HMM analyses. Only high and medium quality MAGs (>50% complete and < 10% contaminated) were included in genome-centric analyses, and the entire dataset (binned and uninned contigs) was considered in gene-centric analyses. For phylogenetic trees, sequences were aligned with muscle v3.8.31 (31), alignment columns were stripped with trimAl v1.4.rev22 (32) using the option -gappyout, and trees were built with FastTree v2.1.10 (33)or UBCG v3.0 (34) and visualized with iToL v6 (35). For calculating average amino acid identity (AAI) between selected MAGs, genomes were gene-called with Prodigal v2.6.3 (36), and amino acid fasta files were used as input to the Kostas Lab online tool (http://enve-omics.ce.gatech.edu/g-matrix/index, (37)). The Genome-to-Genome Distance Calculator v3.0 tool was used online (https://ggdc.dsmz.de/ggdc.php#, (38)). Heat maps were constructed with the package heatmap.2 on RStudio v1.3.959, R v4.0.4.

Metatranscriptomic reads were quality trimmed with Sickle v1.33 (39) using the using the sickle se (single end) or pe (paired-end) options for sanger sequencing (-t sanger). Trimmed transcripts were mapped against the annotated metagenome with Bowtie2 (40) (bowtie2 -D 10 -R 2 -N 1 -L 22 -i S,0,2.50 -q -a -p 30), allowing only one mismatch. Index stats files were imported into RStudio to calculate Transcripts per Million (TPM) values according to the formula TPM = (number of reads / (gene length/10^3^))/ 10^6^), which were used as unit of gene transcription. Aware of differing extraction and sequencing methods for T1 and T2, these two data points have only been used in separate analyses. TPM values were used for bubble plot generation with the packages ggplot on RStudio. All figures were edited on Adobe Illustrator CC v22.1.

### Reverse-transcription quantitative polymerase chain reaction (RT-qPCR)

Selected RNA samples were used for RT-qPCR (Figure 1) in order to confirm patterns emerging from metatranscriptomic analyses. Three primer pairs were selected to quantify transcription of genes of interest: hydrazine synthase subunit A, with *hzsA*-F (5’-WTCGGRTATCARTATGTAG-3’) and *hzsA*-R (5’-AAATGGYGAATCATARTGGC-3’), adapted from previously published primers (41); particulate methane monooxygenase subunit A, with *pmoA*-F (5’-SCGRGTRMAGCCSGGTGAGA-3’) and *pmoA*-R (5’-YGATGGYCCMGGYACMGAGT-3’), designed for this study; and methyl-coenzyme M reductase subunit A, with *mcrA*-F (5’-AAAGTGCGGAGCAGCAATCACC-3’) and *mcrA*-R (5’-TCGTCCCATTCCTGCTGCATTGC-3’) (42). Bacterial and archaeal 16S rRNA gene-targeting primers 16S-F (5’-AAACTYAAAKGAATTGRCGG’) and 16S-R (5’-ACGGGCGGTGWGTRC-3’) were used as housekeeping genes for data normalization, adapted from previously published primers (43). For each sample, 50 ng of RNA was used for the reverse transcription reaction using the RevertAid H Minus First Strand cDNA Synthesis Kit (Thermo Fisher Scientific, Waltham, MA, USA) following manufacturer’s instructions. Each qPCR reaction consisted of 12 μL SYBR Green FastMix (QuantaBio, MA, USA), 1 μL forward and 1 μL reverse primer (10 μM working solutions), 10 μL DEPC-treated water (Thermo Fisher Scientific, Waltham, MA, USA) and 1 μL template single stranded cDNA in a total volume of 25 μL. The qPCR program consisted of the following steps: initial denaturation (94 °C for 5 min), denaturation (94°C for 30 sec), annealing (temperature as below for 30 sec), elongation (72°C for 30 sec), and melting curve (50-95°C, 0.5°C increase per 5 sec). Denaturation, annealing and elongation steps were repeated for 40 cycles. Annealing temperatures for *hzsA*, *pmoA*, *mcrA* and general 16S rRNA gene primers were 55°C, 62°C, 62°C and 55°C, respectively. qPCR performed directly on RNA samples discarded DNA contamination given that all Ct values were above the threshold of 30 cycles. RT-qPCR units were expressed in 2^−ΔΔCT^ by subtracting, first, normalizer with target functional gene CT values, and second, R0 and R3 from R1 and R4 CT values, respectively. Replicates were discarded if CT values above 30.

### Data Availability

Metagenomic trimmed reads, metatranscriptomic trimmed reads, metagenome-assembled genomes and unbinned contigs have been deposited on NCBI under BioProject number PRJNA758578.

## Results

### Diverse microbial community members cycled methane, nitrate, nitrite, ammonium, and sulfide

During the ammonium removal experiment, the bioreactor received the same daily quantities of nitrate (3 mmol), methane (488 mmol) and sulfide (0.67 mmol) as previously described (19). A comprehensive investigation of daily substrate consumption was carried out in the aforementioned study (corresponding the zero time point in Figure 1), which showed that sulfide was completely consumed with a nitrate removal rate of 2.6 mmol per day. The residual nitrate concentrations fluctuated around 4 mM, while no nitrite could be detected in the bioreactor (19). Ammonium was removed from the medium (T1), and nitrate, nitrite and ammonium substrate consumption was monitored for a period of 12 days prior to T2 sample collection (10 weeks without ammonium). Nitrate had a residual concentration of 3 mM, while ammonium, nitrite and sulfide concentrations remained below the detection limit. The overall nitrate and nitrite concentrations were consistent with the previously determined concentrations.

During the sulfide and nitric oxide (NO) toxicity stress experiment, methane, nitrate and ammonium were not provided to the reactor. While sulfide concentrations remained below detection limit for the entire experiment, indicating continuous complete sulfide removal, nitrite accumulated to 870 μM during the second week of the bioreactor receiving only sulfide and nitric oxide, and remained between 100-400 μM during the remainder of the experiment. This indicated that the community removed residual nitrate via denitrification and DNRA for several weeks. Biomass decay was evident from the visual increase in reactor turbidity and, to a degree, change of granule color from red to black, until the end of the experiment.

The composition of the bioreactor microbial community was investigated via a time-series of metagenomic sequencing. In a previous study with this bioreactor, community members included several proteobacteria, *Ca.* Kuenenia stuttgartiensis, *Ca.* Scalindua brodae, *Ca.* Methanoperedens nitroreducens, *Ca.* Methylomirabilis species, and a novel bacterium within the Nitrospirota phylum, *Ca.* Nitrobium versatile, potentially linking sulfur and nitrogen cycling (19, 44–46). Co-assembly of eight samples resulted in 30 high (>90% complete, <5% contaminated) and 29 medium (>50% complete, <10% contaminated) quality metagenome-assembled genomes (MAGs) (Figure 2). We will first introduce the most dominant members and, in the next section, we will discuss their change in abundance over time. The only archaeon detected in the bioreactor was *Ca.* Methanoperedens nitroreducens (MAG 36), a nitrate-reducing anaerobic methanotroph. Bacteria were much more diverse and represented by five MAGs affiliated to candidate phyla (AABM5-125-24, ARS69, FEN-1099, GWC2-55-46, OLB16), two to *Acidobacteriota*, one to *Actinobacteriota*, one to *Armatimonadota*, eleven to *Bacteroidota*, six to *Chloroflexota*, one to *Cyanobacteria*, two to *Methylomirabilota*, two to *Myxococcota*, one to *Nitrospirota*, one to *Omnitrophota*, five to *Planctomycetota*, and twenty to *Proteobacteria*.

**Figure 2.**
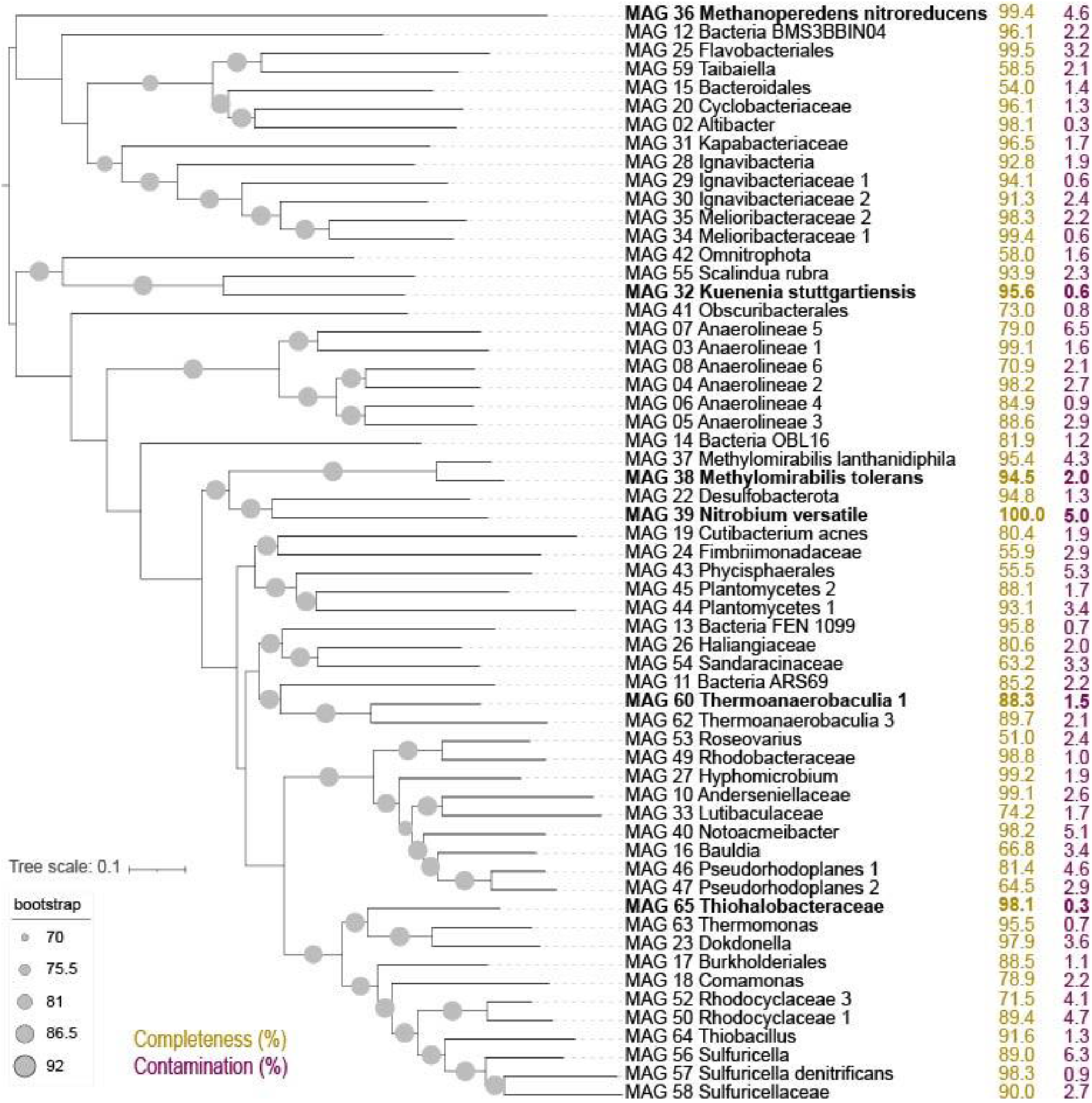
Up-to-date bacterial core gene (UBCG) tree of 92 concatenated genes extracted from high and medium quality MAGs in this study. Genome completeness (in yellow) and contamination (in purple) values in percent are indicated to the right. MAGs in bold were selected from detailed characterization due to large shifts in abundance across experiments.

Two of the *Planctomycetota* MAGs were the anammox bacteria *Candidatus* Kuenenia stuttgartiensis (MAG 32) and *Candidatus* Scalindua rubra (MAG 55). The one *Nitrospirota* MAG 39 was *Candidatus* Nitrobium versatile, which we further characterized in this study. The two MAGs affiliated to *Methylomirabilota* were classified by GTDB-Tk as *Candidatus* Methylomirabilis oxyfera (MAG 37) and *Candidatus* Methylomirabilis sp002634395 (MAG 38). However, phylogenetic and average amino acid identity (AAI) analyses revealed that the first MAG represented a species of *Candidatus* Methylomirabilis lanthanidiphila (100% AAI to the type genome (47)), and the latter was a novel species sharing only 81% AAI to *Candidatus* Methylomirabilis limnetica (Supplemental Figure 1 and 2). The genome-to-genome distance between MAG 38 and *Ca.* M. limnetica ranged from 0.15-0.59, with a probability of DNA-DNA hybridization > 70% ranging from 0-0.08%, supporting AAI results. We have named this novel species *Candidatus* Methylomirabilis tolerans due to its persistence over the sulfide and NO stress experiment (Figure 3). Genome analyses indicated that *Candidatus* Methylomirabilis tolerans had a similar metabolic potential for nitric oxide dismutation coupled to methane oxidation via intracellular oxygen generation as previously described, sharing several other characteristics with *Ca.* Methylomirabilis lanthanidiphila (Table 1). Finally, several previously identified and novel putative sulfide oxidizers were identified based on key gene analyses (Figure 4 and 5): MAG 58 Sulfuricellaceae, MAG 10 Anderseniellaceae, MAG 23 Dokdonella, MAG 40 Notoacmeibacter, MAG 50 Rhodocyclaceae 1, MAG 52 Rhodocyclaceae 3, MAG 57 Sulfuricella denitrificans, MAG 56 Sulfuricella, MAG 64 Thiobacillus, MAG 65 Thiohalobacteraceae, and MAG 63 Thermomonas.

**Table 1.**
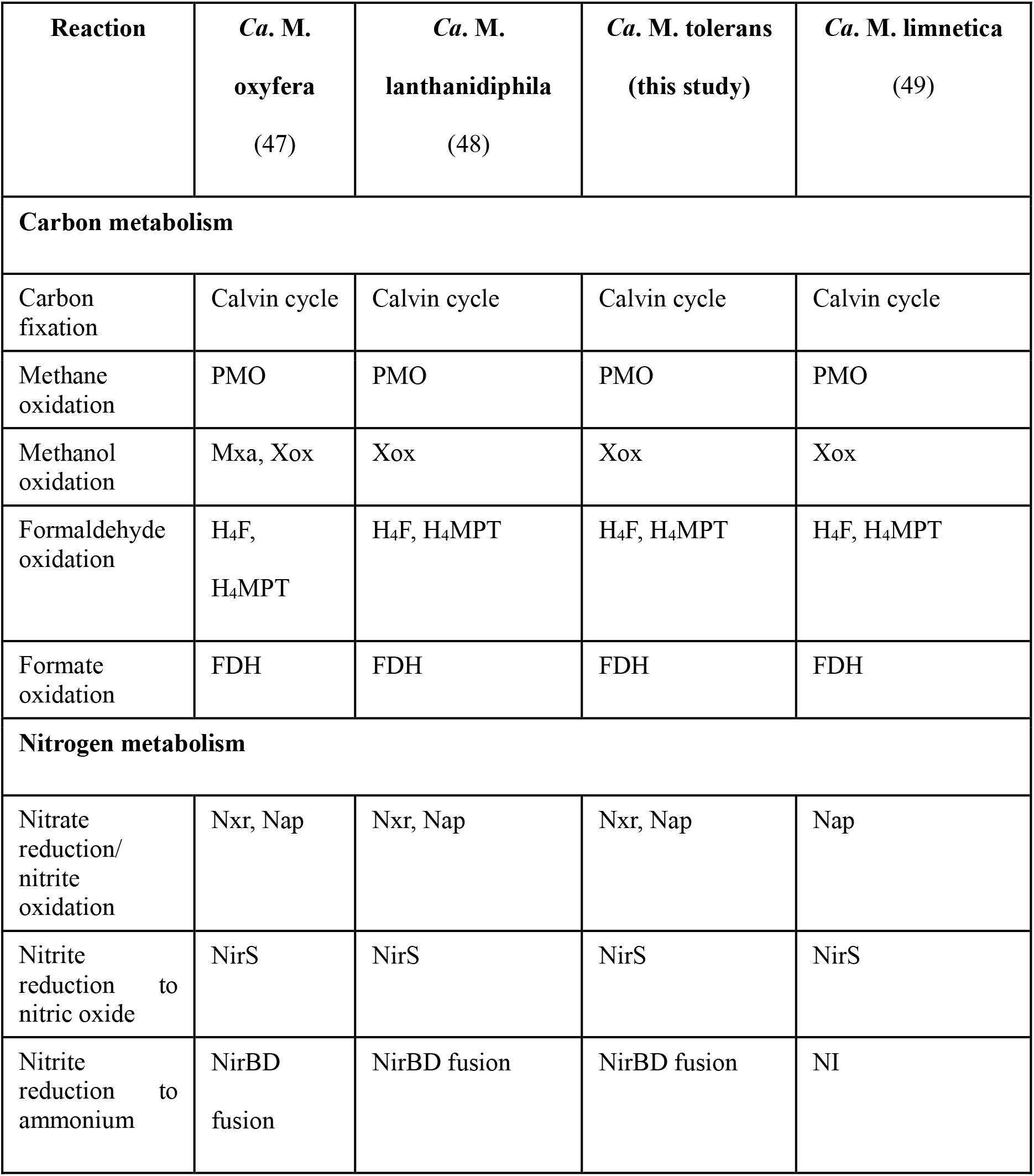
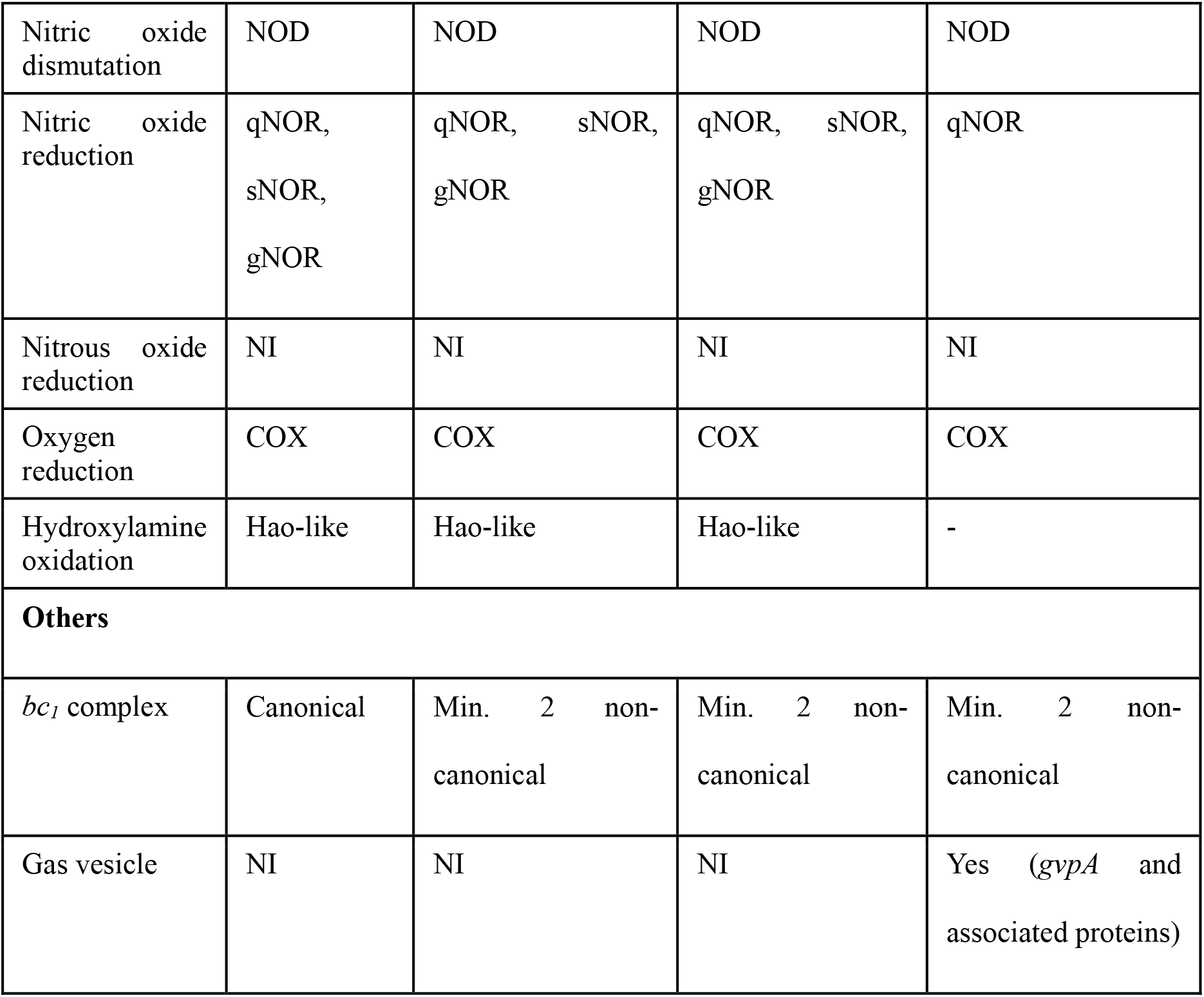
Comparison of functional genes and pathways in selected *Ca.* Methyloramibilis species from this and other studies. Abbreviations are as follows: PMO, particulate methane monooxygenase; Mxa, calcium-dependent methanol dehydrogenase; Xox, lanthanide-dependent methanol dehydrogenase; H_4_F, tetrahydrofolate pathway; H_4_MPT, tetrahydromethanopterin pathway; FDH, formate dehydrogenase; Nxr, membrane-bound nitrate reductase/ nitrite oxidoreductase phylogenetically closest to nitrite-oxidizing enzyme sequences; Nap; periplasmic nitrate reductase; NirS, cytochrome *cd_1_*-containing nitrite reductase; NirBD, cytoplasmic ammonium-forming nitrite reductase; NOD, nitric oxide dismutase; qNOR, quinone-oxidizing electrogenic nitric oxide reductase; sNOR, cytochrome *c*-oxidizing electrogenic nitric oxide reductase; gNOR quinone-oxidizing non-electrogenic nitric oxide reductase; COX, cytochrome *c* oxidase complex; Hao; hydroxylamine oxidoreductase; NI, not identified.

**Figure 3.**
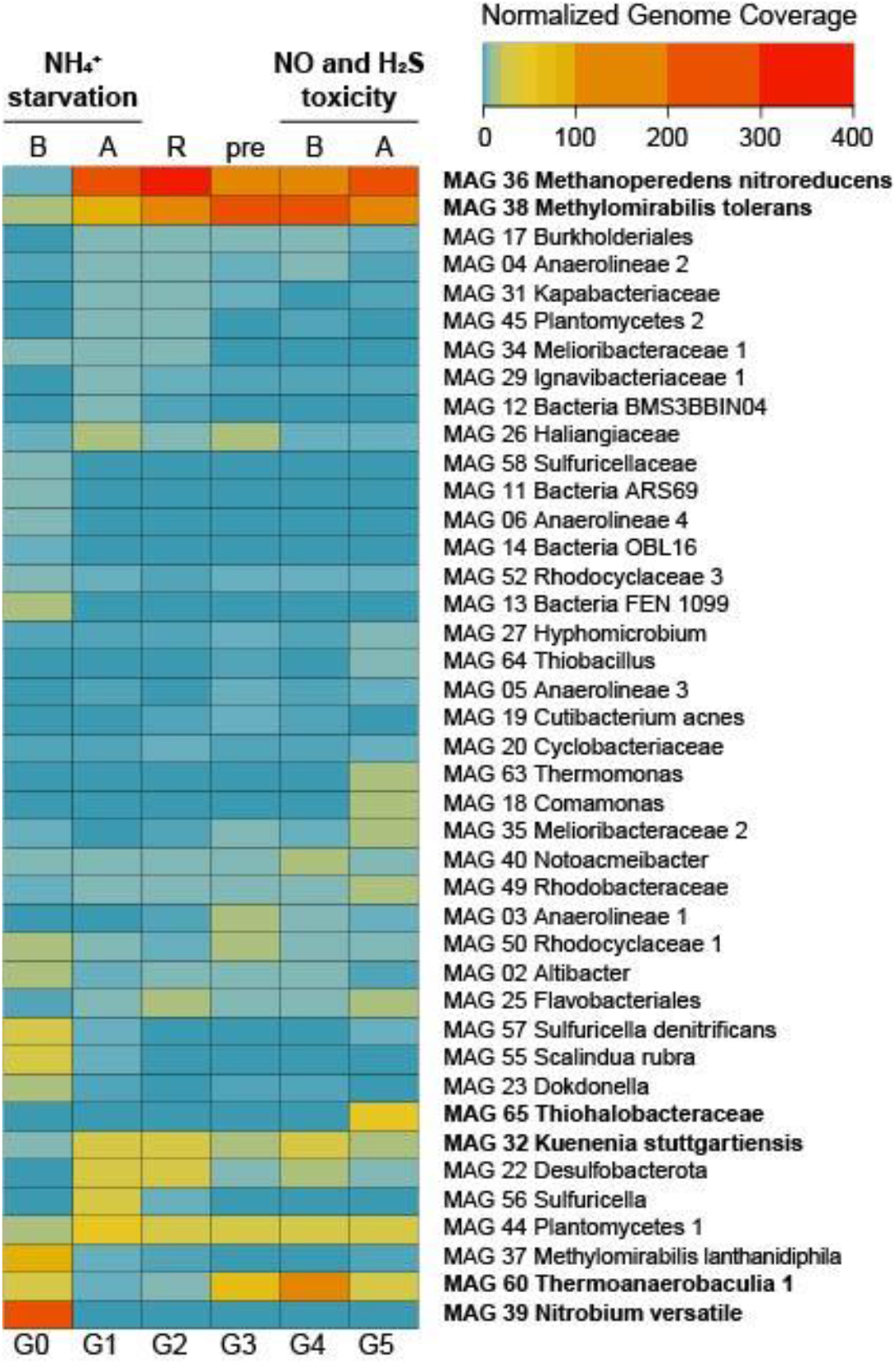
Heat map displaying normalized genome coverages of most abundant MAGs across experimental conditions. Abbreviations are as follows: B, before; A, after; R, recovery; pre, preparatory phase. Experiments are indicated in bold on top. MAGs are ordered based on normalized genome coverage values and abundance change patterns.

**Figure 4.**
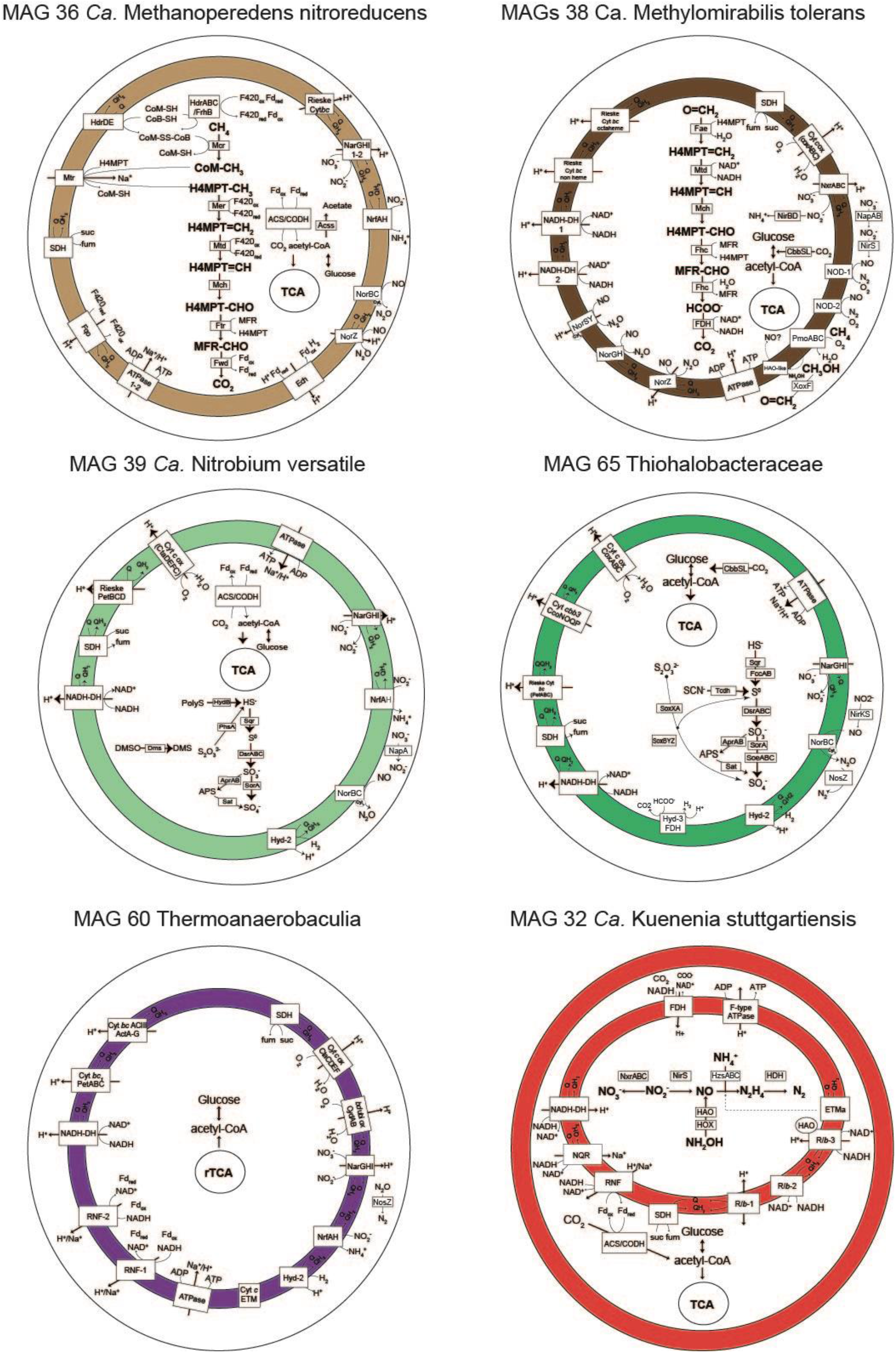
Metabolic reconstruction of six genomes of interest. Only genes with TPM > 0 in metatranscriptomic analyses are shown in selected time points in which the highest number of genes were transcribed (T5 for MAGs 36, 60 and 65, T4 for MAG 38, and T0 for MAGs 32 and 39). Gene abbreviations, loci and annotations are provided in Supplemental Table 4. The gene *nrfH* was absent in MAG 39 *Ca.* Nitrobium versatile (in grey) but it was previously identified in the first MAG representing this organism (19). In MAG 32 *Ca.* Kuenenia stuttgartiensis, the internal compartment represents the anammoxosome.

**Figure 5.**
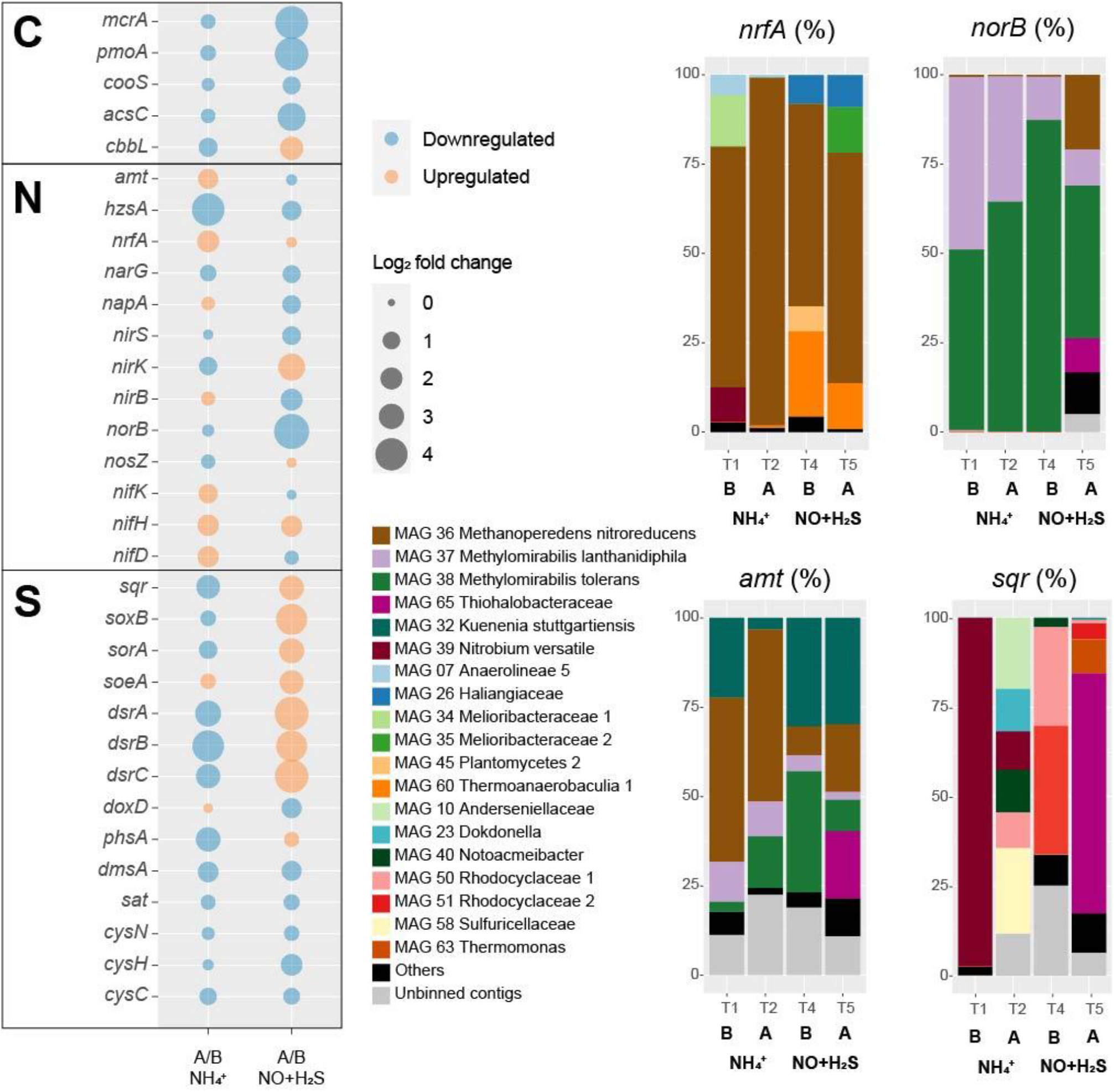
Transcriptional responses to ammonium removal and sulfide and NO toxicity experiments. Log2 fold change, as well as up- or downregulation of genes encoding proteins involved in carbon (C), sulfur (S) and nitrogen (N) cycling, are provide to the right, while taxonomic contributions to selected gene transcription are provided to the left. Under *norB*, subunits of nitric oxide dismutases are also included. Abbreviations are as follows: B, before; A, after.

### MAG coverages revealed shifts in microbial community structure during experimental conditions

Normalized genome coverage for each MAG was used as a proxy for organism abundances across time points. Under regular operation conditions (G0), the most abundant microorganism in the reactor was MAG 39 *Ca*. Nitrobium versatile, with genome coverage of approximately 245x (Figure 3). MAG 37 *Ca.* Methylomirabilis lanthanidiphila had normalized genome coverage of 89x, MAG 60 Thermoanaerobaculia 1 of 26x, MAG 57 Sulfuricella denitrificans of 26x, and MAG 55 Scalindua rubra of 25x. However, large shifts in microbial community structure occurred after ammonium was removed from the medium and subsequently re-introduced (G1). MAG 36 Methanoperedens nitroreducens, which had previously had a normalized genome coverage of 5x, increased to 278x, and MAG 38 Methylomirabilis tolerans, which had a coverage of 19x before, increased to 84x. Other abundant genomes at G1 were MAG 44 Plantomycetes 1 (50x), MAG 22 Desulfobacterota (39x), MAG 32 Kuenenia stuttgartiensis (37x), and MAG 56 Sulfuricella (37x). These genomes remained the most abundant in the bioreactor during the recovery period (G2), with MAG 36 Methanoperedens nitroreducens reaching 366x coverage, and MAG 38 Methylomirabilis tolerans, 110x, indicating that the community reached another distinct stable structure. During the preparatory phase for the sulfide and NO toxicity experiment (G3), MAG 38 Methylomirabilis tolerans became dominant (292x), followed by MAG 36 Methanoperedens nitroreducens (126x), MAG 60 Thermoanaerobaculia 1 (71x), MAG 44 Plantomycetes 1 (33x), and MAG 32 Kuenenia stuttgartiensis (18x). These MAGs remained, in this order, the most abundant genomes in the reactor until the sulfide and NO toxicity experiment started (G4). Finally, at the end of this experiment (G5), shifts in community structure were detected: MAG 36 Methanoperedens nitroreducens returned as the dominant genome (207x), followed by MAG 38 Methylomirabilis tolerans (198x), MAG 65 Thiohalobacteraceae (52x), MAG 44 Plantomycetes 1 (36x), MAG 60 Thermoanaerobaculia 1 (29x), and MAG 32 Kuenenia stuttgartiensis (19x).

### Metatranscriptomic analyses reveal active metabolic pathways in key microbial community members cycling methane, nitrogen and sulfur

In order to investigate transcriptional responses of the bioreactor microbial community to experimental conditions and to unravel microbial pathways likely utilized for metabolic activity in the reactor, we conducted a time series of metatranscriptomic sequencing, from which we calculated transcript per million (TPM) values as unit of gene transcription. We had particular interest in microorganisms that achieved highest abundances or persisted across experimental time points, and selected several key species for in-depth metabolic reconstruction: two key putative sulfur-oxidizing microorganisms (MAG 39 Nitrobium versatile and MAG 65 Thiohalobacteraceae), two key methane oxidizers (MAG 36 Methanoperedens nitroreducens and MAG 38 Methylomirabilis tolerans), and two key nitrogen-cycling microorganisms - the ammonium oxidizer *Ca.* Kuenenia stuttgartiensis (MAG 32) and a putative denitrifying acidobacterium (MAG 60 Thermoanaerobaculia 1).

The *Nitrospirota* MAG 39 Nitrobium versatile, which was the most abundant microbial community member based on genome coverage before the ammonium removal experiment (G0, Figure 3), was estimated to be a 100% complete genome with 5% contamination (Figure 2). Based on metatranscriptomic analyses, we infer that the most likely metabolism performed by this organism was sulfide oxidation coupled to denitrification (Figure 4). This organism was severely affected by the ammonium removal experiment, as indicated by genome coverage (Figure 4). TPM values suggest that *Ca.* Nitrobium versatile became a rare community member but remained transcriptionally active across all time points. When the reactor was primed for the sulfide and NO toxicity experiment (T4), transcripts for 161 genes were detected, including *norC* and *phsA*, while by the end of the experiment (T5), for 130 genes (Supplemental Table 2).

A previously unidentified sulfide:quinone oxidoreductase *sqr* gene in *Ca.* Nitrobium versatile was one of the mostly highly transcribed functional genes (TPM = 0.2±0.05) of this organism in a thriving period (T0, Supplemental Table 2), along with cytochrome *c*-oxidizing nitric oxide reductase genes *norBC* (respectively, TPM = 1.9±0.7 and 2.8±1.2). Sulfide could thus be oxidized to elemental sulfur, which might be the substrate for a dissimilatory sulfite reductase (*dsrABC*) operating in the oxidative direction, despite the two *dsrA* copies in the genome being phylogenetically related to reductive *dsrA* genes (Supplemental Figure 3) and the presence and transcription of a *dsrD* gene. Interestingly, both *dsrABC* copies in the genome were transcribed, one approximately 3 times more than the other (Supplemental Table 2, average TPM 0.3-0.4 vs 0.1). Sulfite could be oxidized to sulfate via transcribed adenosine-phosphosulfate (APS) reductase (*aprAB*) and sulfate adenylyltransferase (*sat*) or via a sulfite:cytochrome *c* oxidoreductase (*sorAB*). Although we did not provide reduced sulfur compounds to the reactor other than sulfide, two genes encoding putative sulfide-generating enzymes were identified and transcribed: a sulfhydrogenase polysulfide reductase (*hydB* subunit beta only, TPM = 0.1±0.06), and a thiosulfate reductase / polysulfide reductase (chain A only, *phsA*, present twice in the genome, TPMs = 0.09±0.05 and 0.016±0.018).

Interestingly, in MAG 39 Nitrobium versatile, a nickel-dependent hydrogenase-(*hyd-2*) and maturation protein-encoding genes were also transcribed at lower levels, indicating that hydrogen could be oxidized. Several other functional genes were transcribed at low levels: membrane-bound and periplasmic nitrate reductase genes (*narGHI* and *napA*), ammonium-forming cytochrome *c_552_*-nitrite reductase (*nrfA*), and anaerobic dimethyl sulfoxide reductase (*dmsABC*). Although previously identified (19), *napB* and *nirBD* were absent in the genome, and *nrfH* was present in the genome but no transcription was detected. All genes encoding subunits of electron transport chain proteins were transcribed (Figure 4). Two genes encoding complexes with homology *caa_3_*-type low affinity cytochrome *c* oxidase proteins in *Acidobacteria* were identified: (i) *ctaCFED*, which was followed by a downstream *sco* assembly protein-encoding gene and a cytochrome *c_6_* (homolog to *petJ*, K08906), which we have putatively denominated *ctaX*, for it could be part of the complex, and (ii) *sco* followed by a downstream *ctaDEFC*. All these subunits were transcribed, except for *ctaF* of the latter complex. A Rieske *bc1* complex was encoded by *petBCD*, which was preceded by a cytochrome *c* protein-encoding gene immediately upstream. Two other copies of *petBC* were present in the genome. All these subunits were transcribed except for the first *petB* (Supplemental Table 4). Finally, the MAG 39 *Ca*. Nitrobium versatile had two ammonium transporter-encoding genes (*amt*), of which only the second was transcribed, solely in T0 (Supplemental Table 4).

The gammaproteobacterial MAG 65 Thiohalobacteraceae, representing an organism that seemed enriched over the sulfide and NO toxicity experiment (Figure 3), was estimated to be a 98.1% complete genome with 0.3% contamination (Figure 2). Based on metatranscriptomic analyses, we infer that the most likely metabolism performed by this organism was sulfide oxidation coupled to denitrification (Figure 4). A sulfide:quinone oxidoreductase-encoding *sqr* gene was among the functional genes with highest transcription by the end of the sulfide and NO toxicity experiment (T5; TPM = 0.43±0.36; Supplemental Table 2, Supplemental Table 4). Also highly transcribed were the genes *dsrA* (TPM = 1.07±0.78), *dsrB* (TPM = 0.95±0.5), *dsrC* (TPM = 4.3±1.1), as well as additional *dsrABC* copies. Sulfite oxidation to sulfate could proceed via proteins encoded by *sorA*, *soeABC* (quinone-sulfite dehydrogenase), as well as *aprAB* and *sat*, which were all transcribed. Sulfur oxidation system *soxBAZYX* genes for thiosulfate oxidation and a thiocyanate dehydrogenase *tcdh* gene were transcribed, indicating that these substrates could also be oxidized, although they were not provided to the reactor. No *soxC* gene was detected in the genome. A cytochrome *cd1*-nitrite reductase *nirS* gene was highly transcribed (TPM = 1.74±0.78), along with all other genes in the denitrification pathway, including a copper-containing nitrite reductase *nirK* gene. Ribulose-bisphosphate carboxylase *cbbSL* genes were among the most highly transcribed in the genome, and all genes encoding subunits of the electron transport chain were transcribed.

MAG 36 (*Ca.* Methanoperedens nitroreducens) was 99.4% complete with 4.6% contamination, and had all genes in the reverse methanogenesis pathway (Figure 4, Supplemental Table 4) as well as potential for carbon fixation via the Wood–Ljungdahl pathway with carbon monoxide dehydrogenase/ acetyl-CoA synthase (CODH/ACS)-encoding genes and acetate production or assimilation via an acetyl-CoA synthetase-encoding *acss* gene. All these genes were transcribed by the end of the sulfide and NO toxicity experiment (T5), as well as electron transport chain-encoding genes (Figure 4). The succinate dehydrogenase *sdhC* subunit was not present in the genome. Remarkably, the genome had genes encoding two copies of the membrane-bound nitrate reductase (*narGHI*), an ammonium-forming nitrite reductase (*nrfAH*), a non-electrogenic cytochrome *c*-oxidizing (cNOR) nitric oxide reductase (*norBC*), and an electrogenic quinone-oxidizing (qNOR) nitric oxide reductase (*norZ*). Methyl-coenzyme M reductase genes *mcrBDGA* had TPM values around ~200-400 before the sulfide and NO toxicity experiment (T4), and ~20-40 after (T5, Supplemental Table 2), indicating that, although *Ca.* Methanoperedens nitroreducens was still abundant by the end of the experiment (Figure 3), it suffered a degree of inhibition. Three hypothetical genes had highest transcription at T5 and were upregulated relative to T4 (Supplemental Table 2).

A second key methane oxidizer was represented by MAG 38 (*Ca*. Methylomirabilis tolerans), 94.5% complete with 2% contamination. This genome had all genes for carbon fixation via the Calvin cycle and for the methane oxidation via particulate methane monooxygenase (*pmoABC*) coupled to nitric oxide dismutation, with four putative *nod* genes present and transcribed. Additionally, nitrate, nitrite and nitric oxide reductase-encoding genes were present and transcribed: *narGHI*, ammonium-forming cytoplasmic NADH-nitrite reductase *nirBD* genes, electrogenic cytochrome *c*-oxidizing (sNOR) nitric oxide reductase-encoding *norSY* genes, non-electrogenic quinone-oxidizing (gNOR) nitric oxide reductase-encoding *norGH* genes, and *norZ*. All four genes encoding the two subunits of the nitric oxide dismutases NOD-1 and NOD-2 had some of the highest transcription (TPM ~7-60) when the community was being primed for the sulfide and NO toxicity experiment, receiving ~1-5% external NO in the reactor along with methane, ammonium and sulfide (T4), as well as *pmoABC* (TPM ~6-22) and lanthanide-dependent methanol dehydrogenase-encoding *xoxF* gene (TPM = 5.5± 3.5) (Supplemental Table 2). Although *Ca*. Methylomirabilis tolerans was still abundant after the sulfide and NO toxicity experiment (Figure 3), TPM values for all aforementioned functional genes decreased 10-100 fold from T4 to T5, indicating that the microorganism suffered a degree of inhibition (Supplemental Table 2).

MAG 60 Thermoanaerobaculia 1, 88.3% complete with 1.5% contamination, had highest normalized genome coverage when the reactor was being primed for the sulfide and NO toxicity experiment (G4, Figure 3). Based on metatranscriptomic analyses, the most likely metabolism performed by this organism was heterotrophic denitrification (Figure 4). A nitrous oxide reductase *nosZ* gene was among the functional genes with highest transcription (T4, TPM = 0.21±0.17), followed by *narGHI* and *nrfAH*. Interestingly, hydrogenase-2 and low affinity cytochrome *c* oxidase-encoding *ctaCDEF* genes were also transcribed, as well as all electron transport chain-encoding genes, including two RNF complexes (Supplemental Table 2 and 4).

Finally, *Ca.* Kuenenia stuttgartiensis, represented by MAG 32, 95.6% complete with 0.6% contamination, was one of two microorganisms that persisted throughout all regular and experimental conditions (Figure 3). Interestingly, the transcription of hydrazine synthase *hzsABC* genes decreased 18-27x during the ammonium removal experiment (T1 to T2), and 2-5x during the sulfide and NO toxicity experiment (T4 to T5), suggesting that substrate deprivation was a strong stressor. MAG 32 had all genes encoding substrate oxidation and electron transport chain proteins in the anammox pathway, as well carbon fixation via the Wood–Ljungdahl pathway, which were transcribed at all time points (Figure 4). The second microorganism that persisted throughout all regular and experimental conditions was represented by MAG 44, 93.1% complete with 3.4% contamination, a poorly classified *Planctomycetes* genome. Based on metagenomic and transcriptomic analyses, the most likely metabolism performed by this organism was heterotrophic denitrification via *narGHI* and *nirS* (Supplemental Table 2). While normalized genome coverage of this MAG varied only between 16-50x across regular and experimental conditions, summed TPM values for all genes transcribed in each time point were highest in the preparatory phase for the sulfide and NO toxicity experiment, as well as before and after the experiment (T3-5; TPM = ~168-361) in comparison to previous time points (T0-2; TPM = ~33-43; Supplemental Table 2).

### Microbial community transcriptional responses to substrate removal and toxicity stresses provide insights into community dynamics and resiliency

Gene-centric analyses aimed to elucidate the transcriptional responses of the reactor community to two experimental conditions. The first, ammonium removal from the medium while methane, sulfide and nitrate were still provided to the reactor, tested the strength of microbial interactions for the supply of ammonium via dissimilatory nitrate reduction to ammonium (DNRA) to community members - in particular, to anammox bacteria. The second, sulfide and nitric oxide addition while no other substrates were provided to the reactor, aimed to enrich sulfide-oxidizing denitrifiers and to test the limits of microbial community resiliency to these stresses.

A major response to ammonium removal was a general inhibition of methane, ammonium and sulfide oxidizers, as indicated by downregulation of their key genes *mcrA*, *pmoA*, *hzsA*, *sqr*, *soxB*, *sorA*, and *dsrABC* (Figure 5). RT-qPCR of *mcrA*, *pmoA*, and *hzsA* confirmed these trends (Supplemental Table 3). Not surprisingly, anammox bacteria were particularly affected, with an, approximately, 18-fold decrease in *hzsA* transcription. On the other hand, nitrogen fixation *nifDHK* genes, ammonium transporter *amt* genes, and the ammonium-forming nitrite reductase *nrfA* gene were upregulated, likely to compensate for the ammonium limitation. *Ca.* Methanoperedens nitroreducens accounted for 97.6% of *nrfA* transcription and 48.2% of *amt* transcription 10 weeks after ammonium was completely removed from the medium (T2). Other taxa that shared a large proportion of *amt* transcription in T2 were *Ca.* Methylomirabilis tolerans (14.5%), *Ca.* Methylomirabilis lanthanidiphila (9.6%), and *Ca.* Kuenenia stuttgartiensis (3.2%). The *hzsA* gene of *Ca.* Kuenenia stuttgartiensis had at least 20x higher transcription than *hzsA* of *Ca.* Scalindua rubra across all time points, although *Ca.* Scalindua rubra had higher genome coverage before experiments were conducted (Figure 3). Finally, minor changes included the upregulation of the periplasmic nitrate reductase *napA* gene, cytoplasmic ammonium-forming nitrite reductase *nirB* gene, quinone-sulfite dehydrogenase *soeA* gene, and quinone-thiosulfate dehydrogenase *doxD* gene.

A major response to the sulfide and NO toxicity experiment was upregulation of all sulfur oxidation genes (*sqr*, *soxB*, *sorA*, soeA, and *dsrABC*) (Figure 5). MAG 65 Thiohalobacteraceae accounted for 67% of *sqr* transcription by the end of the experiment (T5). Interestingly, the transcription of nitric oxide reductase *norB* genes decreased 32 times. However, *norB* TPM values were the highest among all nitrogen cycle marker genes across all time points in this study except T5 (average TPM = 5.22), in which *hzsA* TPM values were higher (average TPM = 7). At T5, *Ca.* Methylomirabilis tolerans accounted for 43% of *norB* transcription at T5, which included NOD-1 (18%), NOD-2 (19%), and qNOR (6%)-encoding genes, followed by *Ca.* Methanoperedens nitroreducens (21%, with the qNOR-encoding gene accounting for 13% and cNOR-encoding gene for 8%), *Ca.* Methylomirabilis lanthanidiphila (10%) and MAG 65 Thiohalobacteraceae (9.6%). The transcription of several other genes encoding proteins involved in denitrification (*narG*, *napA*, *nirS*, and *nirB*) was downregulated. However, *nirK* had a 12-fold increase in transcription, and *nosZ* had a minor increase. Methane oxidation marker genes were downregulated, as indicated by a 19-fold decrease in *mcrA* transcription and 21-fold decrease in *pmoA* transcription. Transcription of the anammox marker gene *hzsA* decreased 2.7 fold. RT-qPCR of *mcrA*, *pmoA*, and *hzsA* confirmed these trends (Supplemental table 3). Carbon fixation via CODH/ACS was downregulated, but CO_2_ fixation via ribulose-bisphosphate carboxylase-oxygenase was upregulated.

## Discussion

In this study, we conducted metagenomic and metatranscriptomic analyses in order to unravel metabolic pathways, microbial dynamics, and responses to stresses in a bioreactor community mimicking anoxic, brackish coastal sediment conditions. Microbial interactions and resiliency were tested by removal and addition of key substrates. Removal of ammonium from the medium had major effects on microbial activity and community structure: *Ca.* Methanoperedens nitroreducens dominated dissimilatory nitrate reduction to ammonium (DNRA) activity (Figure 5), and, after ammonium was reintroduced in the medium, became the most abundant community member (Figure 3). *Ca.* Kuenenia stuttgartiensis replaced *Ca.* Scalindua rubra as the most abundant anammox bacteria, several sulfide oxidizers replaced *Ca.* Nitrobium versatile, and *Ca.* Methylomirabilis tolerans dominated over *Ca.* Methylomirabilis lanthanidiphila (Figure 3 and 5). These results suggest strong microbial cooperation between *Ca.* Methanoperedens nitroreducens and *Ca.* Methylomirabilis species for the exchange of nitrite under the abundance of methane. Before the ammonium removal experiment, TPM values indicate that *Ca.* Methanoperedens nitroreducens reduced nitrate to nitrite, which was reduced to nitric oxide mostly by *Ca.* Methylomirabilis lanthanidiphila. Nitric oxide was then shared among three main organisms: *Ca.* Methylomirabilis lanthanidiphila, *Ca.* Methylomirabilis tolerans, and *Ca.* Nitrobium versatile. However, under ammonium limitation, *Ca.* Methanoperedens nitroreducens further reduced nitrite to ammonium via DNRA, and this had a cascade effect that changed microbial community structure, resulting that *Ca.* Methylomirabilis tolerans dominated and *Ca.* Methylomirabilis lanthanidiphila and *Ca.* Nitrobium versatile concomitantly decreased in abundance and activity. These results suggest that *Ca.* Methylomirabilis tolerans was a more competitive organism in scavenging nitric oxide. However, the mechanisms remain to be elucidated. Both *Ca.* Methylomirabilis species encoded lanthanide-dependent methanol dehydrogenases and could be enriched in the bioreactor due to cerium being provided as part of the medium (47, 48, 50).

Anammox bacteria were more strongly limited by ammonium removal than *Ca.* Methylomirabilis species, presumably due to the dual competition for both ammonium and nitrite. Although *Ca.* Methylomirabilis organisms have a lower affinity for nitrite (K_s_ = 7 μM; *Ca.* M. oxyfera and *Ca.* M. lanthanidiphila (50)) than anammox bacteria (K_s_ =0.2-3 μM; *Ca.* K. stuttgartiensis and *Ca.* Scalindua sp. (51)), additional competition for ammonium could have caused anammox to be less fit. Yet, *Ca.* K. stuttgartiensis (K_s_ < 5 μM for ammonium (52)) seemed to better withstand ammonium starvation than *Ca.* Scalindua rubra (K_s_ = 3 μM for ammonium in several *Ca.* Scalindua species (51)), indicating that *Ca*. K. stuttgartiensis’ affinity for ammonium could be higher than that of *Ca*. Scalindua. *Ca.* Nitrobium versatile, on the other hand, seemed to rely on nitric oxide for sulfide oxidation, decreasing in abundance as *Ca.* Methanoperedens nitroreducens (K_s_ = 2.1 ± 0.4mg N L^−1^ for nitrate (53)) and *Ca.* Methylomirabilis tolerans increased (Figure 3). All three organisms had potential to generate ammonium (Figure 4), but *Ca.* Methanoperedens nitroreducens dominated DNRA activity (Figure 5). Therefore, we conclude that DNRA from *Ca.* M. nitroreducens alone could not sustain anammox activity under ammonium, nitrite and organic carbon (except methane) limitation, favoring *Ca.* Methylomirabilis tolerans, which likely outcompeted *Ca.* N. versatile and anammox for nitrite and nitric oxide. These results are contrasting to estuary ecosystems in which anammox activity could be sustained by DNRA, positively correlating to sulfide and sediment organic carbon content (54). High sediment organic carbon to nitrate ratio and ferrous iron availability have been reported to favor DNRA over denitrification in estuary ecosystems (55). Having methane as the only organic carbon source (at saturation) and a high nitrate load (3 mmol/day) could have thus modulated DNRA activity to become insufficient to sustain anammox and changed microbial community structure.

We also tested microbial community resiliency to sulfide and NO toxicity while attempting to enrich sulfide-oxidizing nitric oxide reducers. We found genomic potential and transcriptional evidence for this metabolism in *Ca.* Nitrobium versatile, the dominant microorganism in the bioreactor community before the ammonium removal experiment (Figure 3). However, *Ca.* Nitrobium versatile became a rare community member after this experiment and could no longer be enriched. Instead, MAG 65 Thiohalobacteraceae, likely performing sulfide oxidation coupled to denitrification, increased in abundance (Figure 3, 4 and 5). The sulfide and NO toxicity experiment revealed unusual resiliency of *Ca.* M. nitroreducens and *Ca.* M tolerans, which persisted as the most abundant community members (Figure 3) even though methane and nitrate were no longer provided to the reactor for the 7 weeks of the experiment, consistent with the downregulation of *mcrA* and *pmoA* (Figure 5, Supplemental Table 3). Given that the hydraulic retention time of the reactor was 5 days, it is unlike that our sequencing results and the dominance of *Ca.* M. nitroreducens and *Ca.* M tolerans simply reflect older decaying biomass.

MAG 65 Thiohalobacteraceae, MAG 44 Plantomycetes 1, MAG 60 Thermoanaerobaculia 1, and MAG 32 *Ca*. Kuenenia stuttgartiensis followed as the most abundant community members at the end of the experiment. While the first three could make use of sulfide or organic carbon from decaying biomass as electron donors and residual nitrate or nitrite and nitric oxide as electron acceptors, *Ca.* K. stuttgartiensis could have used ammonium generated from decaying biomass and nitrite from residual nitrate reduction or nitric oxide (56). Of these four MAGs, only MAG 65 Thiohalobacteraceae increased in abundance and had both *nirK* and *norB* transcription increased during the sulfide and NO toxicity experiment (Figure 5, Supplemental Table 2), while the other three suffered a degree of inhibition inferred from both decreases in genome coverage (Figure 3) and marker gene transcriptional activity (Figure 5, Supplemental Table 2). Given that sulfide remained below detection limit (0.15 μM) during all time points, we infer that this inhibition is attributed to nitrite (~100-400 μM) and nitric oxide (~10-13% of headspace). While nitrite and nitric oxide were likely toxic for several community members, MAG 65 Thiohalobacteraceae seemed to have taken advantage of these compounds as terminal electron acceptors.

MAG 65 Thiohalobacteraceae had similar metabolic potential as the gammaproteobacterium *Thiohalabacter thyonaticus*, which has sulfur oxidation genes encoding FccAB, DsrABC, AprAB, Sat, SoeABC, and SoxXABYZ, as well as thiocyanate dehydrogenase and carbon fixation via the Calvin cycle (57). Additionally, MAG 65 had three *sqr* copies and two *sorA* copies. Interestingly, MAG 39 *Ca.* N. versatile also had a sulfur oxidation pathway that included *sqr*, *dsrABC*, *sorA* or *aprAB* and *sat*. Umezawa *et al.* (2020) described the YTD gene cluster composed of genes *yedE*-like, *tusA*, *dsrE*-like, *chp-1* and *chp-2*, which encoded proteins for sulfur disproportionation in *Nitrospirota* (44). As in this previous study, we also detected an impartial YTD gene cluster in *Ca.* N. versatile (MAG 39), indicating that it might lack sulfur disproportionation potential via this cluster, contrasting to the *Nitrospirota* microorganism species 45J (44), which shares 64% average amino acid identity (AAI) to *Ca.* N. versatile. The lack of a complete YTD gene cluster is also described in the *Nitrospirota* microorganism *Candidatus* Sulfobium mesophilum (44, 46), which has *sqr* and *dsrABCD* as well as *napAB*, and *nrfAH*, and shares 57% average amino acid identity to *Ca.* N. versatile. These AAI values indicate that *Ca.* N. versatile and species 45J are likely part of the same *Ca.* Nitrobium genus but distinct species, and that together with *Ca.* Sulfobium mesophilum they form a family-level taxonomic group (58). Umezawa *et al.* (2021) isolated a sulfur-disproportionating microorganism that was named *Dissulfurispira thermophila*, a strain belonging to species 45J (45), therefore matching the genus *Ca.* Nitrobium described by our group in 2017 (19).

*Ca.* Sulfobium mesophilum and *Ca.* N. versatile both have *dsrA* genes that affiliate with bacterial-type reductive sequences, and a *dsrD* gene, suggested as a potential marker for sulfate reduction, absent in sulfur oxidizers that utilize the reverse Dsr pathway (59–61). However, our metatranscriptomic results suggest that *Ca.* N. versatile was performing sulfide oxidation. Therefore, it seems that *Ca.* N. versatile, similarly to the deltaproteobacterium *Desulfurivibrio alkaliphilus*, was yet another example of a microorganism disguised as a sulfate reducer (62). Interestingly, *Ca.* Sulfobium mesophilum and *Desulfurivibrio alkaliphilus* could couple sulfide oxidation to DNRA, a potential also present in MAG 39, with transcripts to *nrfA* detected in our study (T0; TPM = 0.030±0.031). Given that *norBC* had significantly higher transcriptional detection than *nrfA* and other denitrification genes (T0, TPM *norB* = 1.9±0.7, TPM *norC* = 2.8±1.1), it is more likely that *Ca.* N. versatile coupled sulfide oxidation to nitric oxide reduction. Given that *Ca.* N. versatile could not be enriched under sulfide and NO, we hypothesize that MAG 65 Thiohalobacteraceae was a more competitive microorganism. However, the mechanism and substrate affinities remain to be elucidated.

In our study, methanotrophs could withstand prolonged periods of ammonium, methane and nitrate deprivation as well as exposure to elevated concentrations of nitrite and nitric oxide. During these disturbances, *Ca.* Methanoperedens nitroreducens and *Ca.* Methylomirabilis tolerans did not thrive, as indicated by downregulation of *mcrA* and *pmoA*, but tolerated stresses and persisted as abundant community members. These results suggest that methane oxidation could be a relatively stable community function in coastal ecosystems under the stresses investigated in this study. Interestingly, the *Ca.* Methanoperedens nitroreducens genome (MAG 36) encoded nitric oxide reductases (Figure 4), which had not been described before in this organism. The qNOR-encoding gene, present in the same contig as *mcr*, *nar* and *nrf* genes, had increased transcriptional activity by the end of the sulfide and NO stress experiment (Figure 5). Future studies should further evaluate lateral transfer of *nor* genes in *Methanoperendaceaea*, which seem prone to acquire novel metabolic traits via later gene transfer events (63), potentially via recently described borgs (64), as well as metabolic flexibility of *Ca.* Methanoperedens nitroreducens under methane deprivation. Hypothetical genes in *Ca.* Methanoperedens nitroreducens identified in this study with high transcription and upregulation in response to stresses could be targets for these future investigations.

Sulfide oxidizers had differential resiliency to the stresses investigated in this study, dynamically changing in abundance and transcriptional activity across regular and experimental conditions. However, sulfide was completely removed at all time points, likely due to functional redundancy in the microbial community, indicating that sulfide oxidation could also be a relatively stable community function in coastal ecosystems under the investigated stresses. Finally, in our study, denitrification was a dominant nitrogen-cycling pathway, as previously suggested in coastal sediments of the Bothnian Sea (65) - particularly, the nitric oxide reduction step, as indicated by *norB* TPM values (Supplemental Table 2). Anammox activity, as indicated by *hzsA* TPM values, also had a significant contribution to nitrogen cycling, and dominated at the end of the sulfide and nitric oxide stress experiment (T5). This community function was highly impacted by ammonium deprivation, but could be restored when favorable conditions were reestablished. On the other hand, DNRA, as indicated by *nrfA* TPM values, was a minor nitrogen-cycling pathway, but under ammonium deprivation it became more significant. Therefore, anammox and DNRA activity might similarly oscillate in coastal ecosystems under the stresses investigated in this study.

In conclusion, this study contributed to the elucidation of metabolic pathways for carbon, sulfur and nitrogen cycling in a bioreactor community mimicking anoxic, brackish coastal sediment conditions, as well as shifts in microbial abundance and transcriptional activity in response to prolonged substrate deprivation and exposure to toxic compounds. These results will help understanding complex microbial interactions and functions in dynamic costal ecosystems, and should be considered into future modelling efforts that aim to predict coastal ecosystem responses to environmental change.

## Acknowledgements

We thank Dr. Arjan Pol and Dr. Tom Bergen for research discussions and suggestions, and Dr. Michiel in’t Zandt for data visualization inspiration. Additionally, we are grateful to Prof. Guo-Jun Xie and Dr. Nie for sharing *Ca.* Methylomirabilis genomes with us in busy times. This research was funded by the Soehngen Institute of Anaerobic Microbiology Gravitation Grant 024.002.002, European Research Council Synergy Grant MARIX 854088, and Nederlandse Organisatie voor Wetenschappelijk Onderzoek Grant ALWOP.293.

## Supplemental Files

**Supplemental Figure 1.**
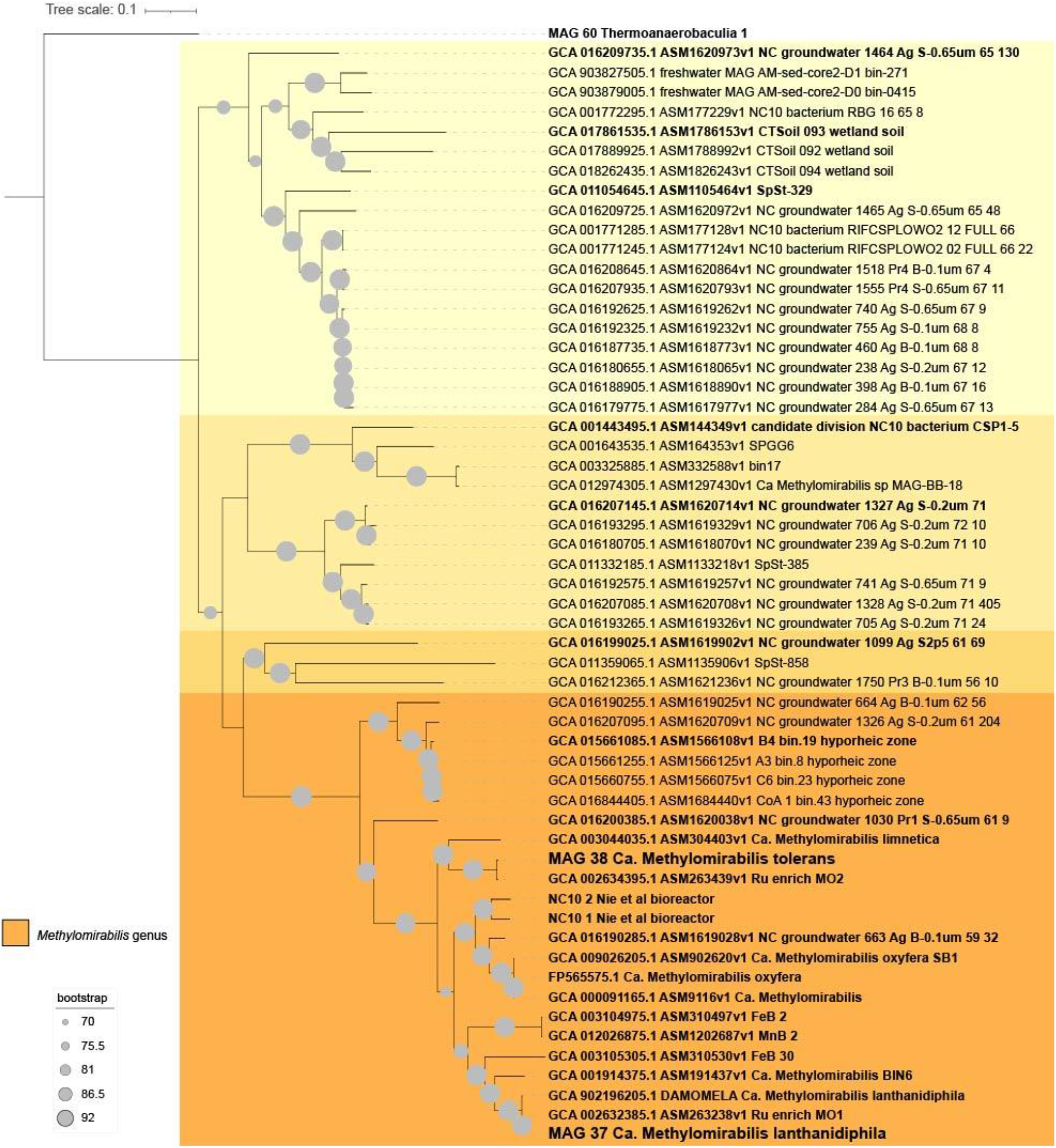
Phylogenetic tree of NC10 phylum-affiliated genomes retrieved from NCBI (accession numbers are indicated), from Nie et al upon request (12), and from this study (MAG 37 and 38). Ninety-two genes were extracted with UBCG, concatenated and aligned with FastTree. The *candidatus* genus Methylomirabilis is highlighted in orange. Genomes in bold were used for average amino acid identity analyses in Supplemental Figure 2.

**Supplemental Figure 2.**
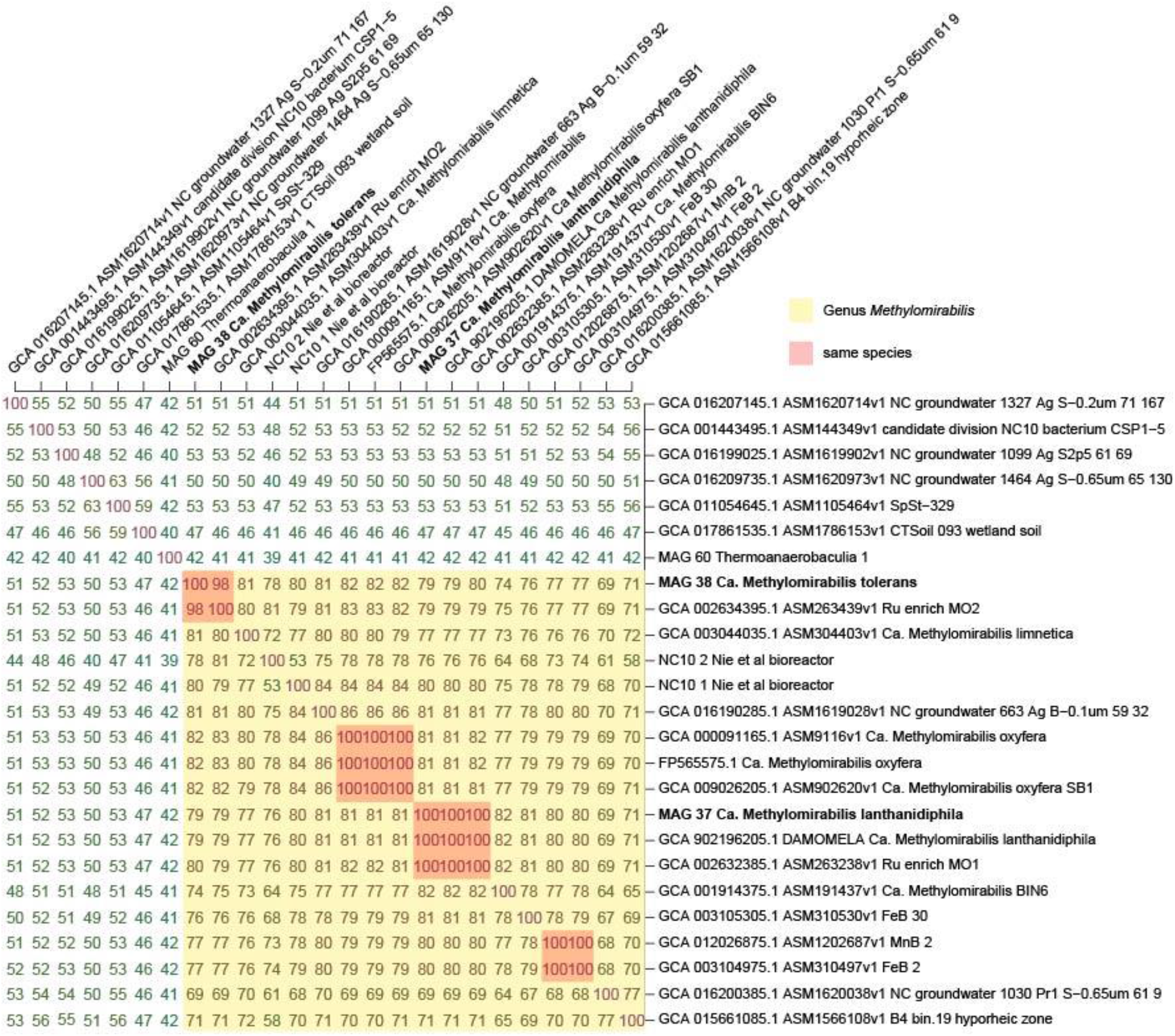
Average amino acid identity (AAI) matrix highlighting the *candidatus* genus Methylomirabilis (yellow) and some genomes of the same species (red).

**Supplemental Figure 3.**
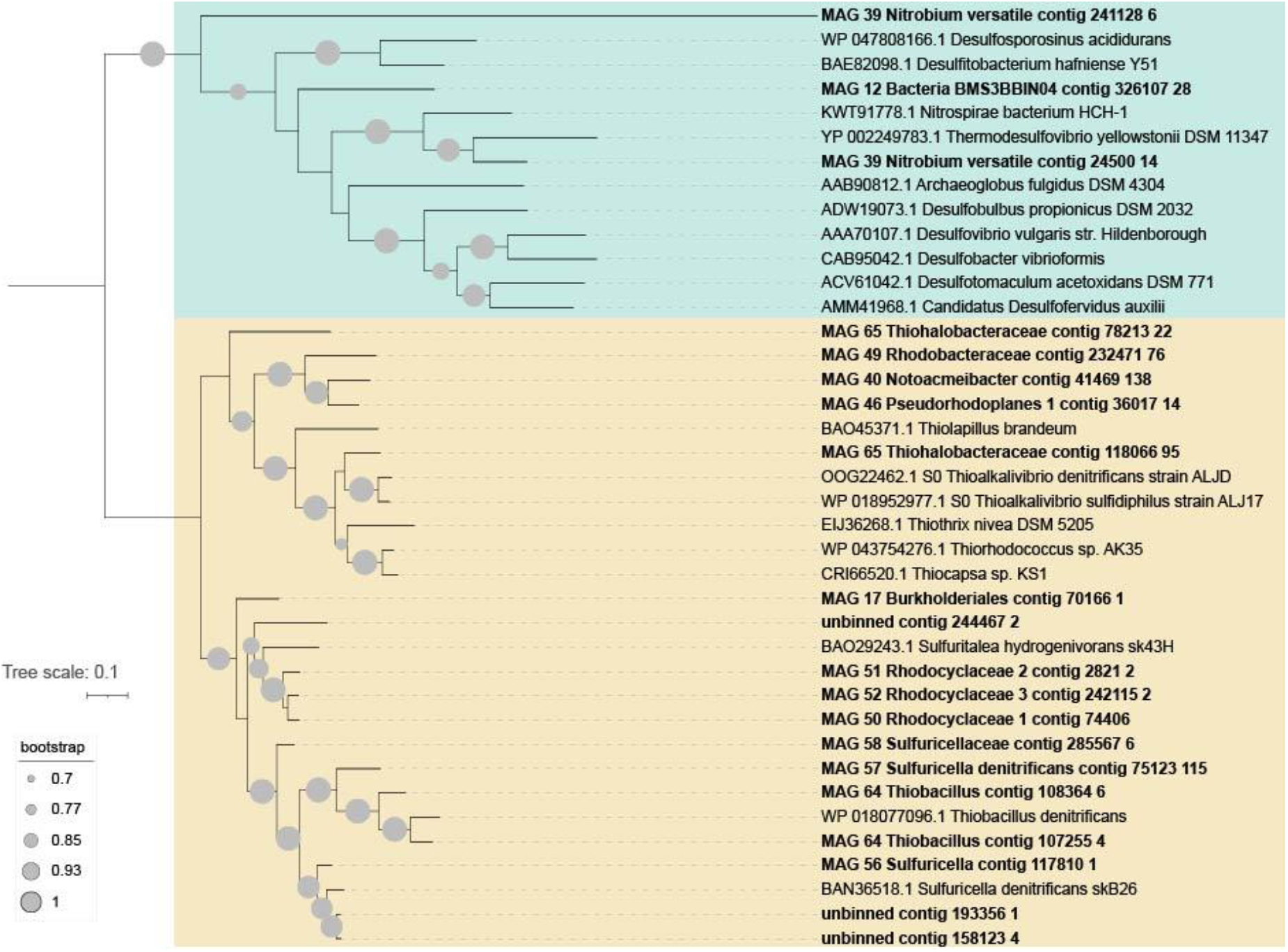
Phylogenetic tree of DsrA sequences from this study (in bold) and reference sequences (indicated by NCBI accession numbers). Sequences putatively assigned to the sulfur reduction direction are highlighted in blue, and to the oxidative direction are highlighted in yellow.

**Supplemental Table 1.**
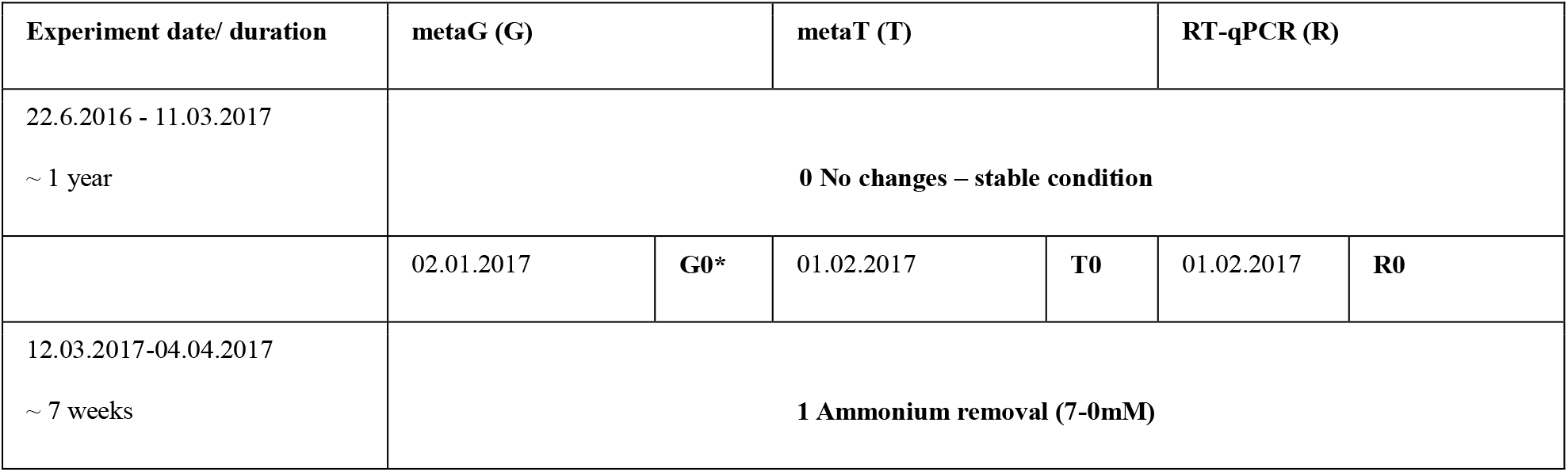
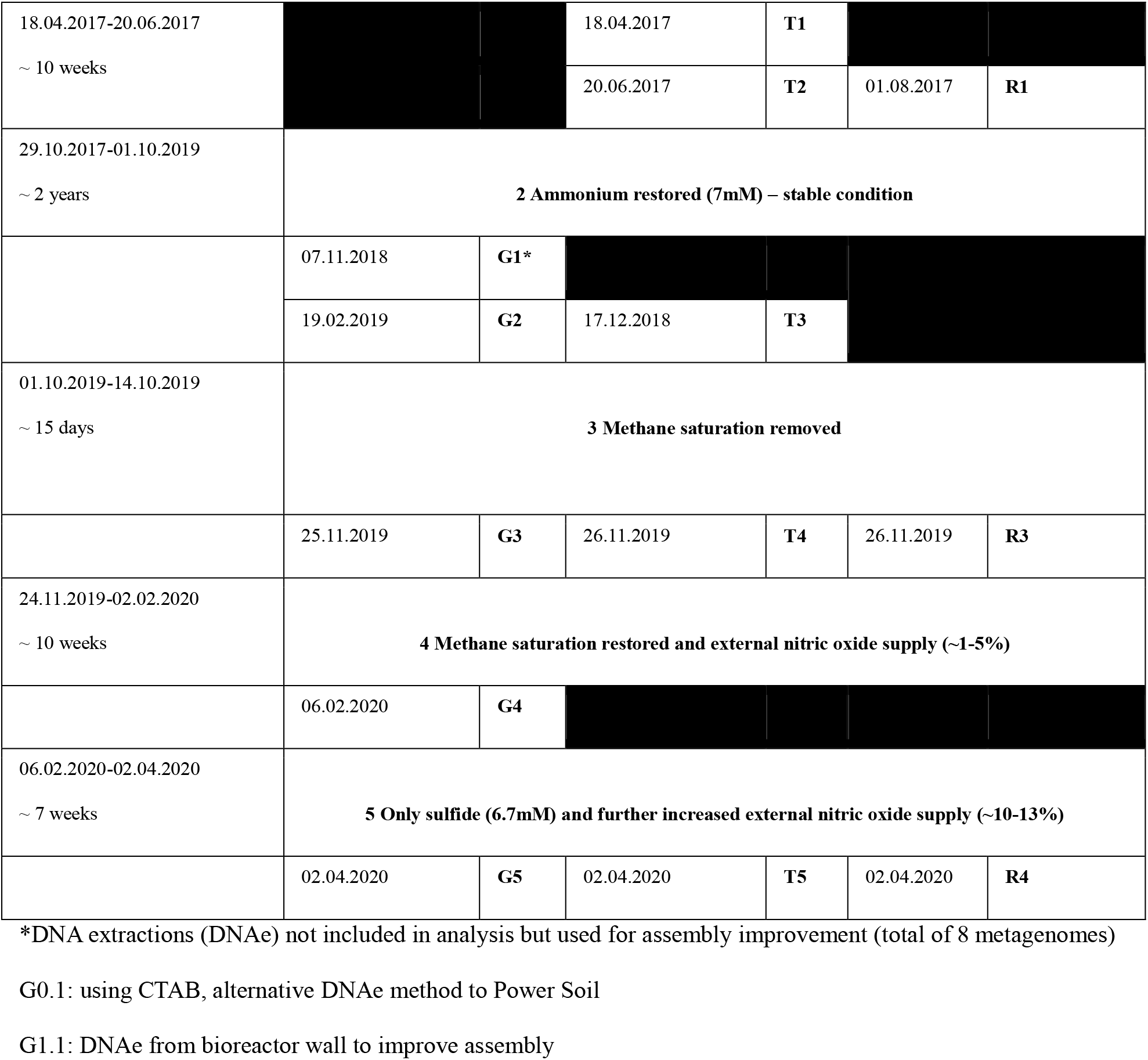
Specific dates, duration (left) and numbering (right) of the ammonium removal and toxicity stresses experiments. Right columns- specific biomass collection dates and numbering for metagenomics (G), metatranscriptomics (T), and RT-qPCR (R) before/after substrate removal and stresses. (dd.mm.yyyy).

**Supplemental Table 2.** Transcript per million (TPM) values for each gene and genome in this study (excel spreadsheet).

**Supplemental Table 3.** RT-qPCR R0-R1 and R3-R3 results calculated in 2^^^ΔΔCT values for selected functional genes using three biological and technical duplicates per time point (R).

| $2^{(-\Delta\Delta\text{CT})}$ | *hzsA* | *pmoA* | *mcrA* |
| --- | --- | --- | --- |
| R0-R1 | 0.00 | 1.24 | 1.15 |
| R3-R4 | 0.02 | 0.00 | 0.00 |

**Supplemental Table 4.** Gene annotations and loci for MAGs included in Figure 4 (excel spreadsheet).

